# Oncogenic hijacking of a developmental transcription factor evokes therapeutic vulnerability for ROS-induction in Ewing sarcoma

**DOI:** 10.1101/578666

**Authors:** Aruna Marchetto, Shunya Ohmura, Martin F. Orth, Jing Li, Fabienne S. Wehweck, Maximilian M. L. Knott, Stefanie Stein, David Saucier, Chiara Arrigoni, Julia S. Gerke, Michaela C. Baldauf, Julian Musa, Marlene Dallmayer, Tilman L. B. Hölting, Matteo Moretti, James F. Amatruda, Laura Romero-Pérez, Florencia Cidre-Aranaz, Thomas Kirchner, Giuseppina Sannino, Thomas G. P. Grünewald

**Affiliations:** Max-Eder Research Group for Pediatric Sarcoma Biology, Institute of Pathology, Faculty of Medicine, LMU Munich, Germany; Institute of Pathology, Faculty of Medicine, LMU Munich, Germany; Department of Pediatrics and Molecular Biology, University of Texas Southwestern Medical Center and Children’s Medical Center, Dallas, TX, USA; Regenerative Medicine Technologies Laboratory, Ente Ospedaliero Cantonale (EOC), Lugano, Switzerland; German Cancer Consortium (DKTK), partner site Munich, Germany; German Cancer Research Center (DKFZ), Heidelberg, Germany

**Keywords:** SOX6, Ewing sarcoma, Elesclomol, ROS, EWSR1-FLI1, TXNIP

## Abstract

Ewing sarcoma (EwS) is an aggressive childhood cancer likely originating from mesenchymal stem cells or osteo-chondrogenic progenitors. It is characterized by fusion oncoproteins involving EWSR1 and variable members of the ETS-family of transcription factors (in 85% FLI1). EWSR1-FLI1 can induce target genes by using GGAA-microsatellites (mSats) as enhancers.

Here, we show that EWSR1-FLI1 hijacks the developmental transcription factor SOX6 – a physiological driver of proliferation of osteo-chondrogenic progenitors – by binding to an intronic GGAA-mSat, which promotes EwS growth *in vitro* and *in vivo*. Through integration of transcriptome-profiling, published drug-screening data, and functional *in vitro* and *in vivo* experiments, we discovered that SOX6 interferes with the antioxidant system resulting in constitutively elevated reactive oxygen species (ROS) levels that create a therapeutic vulnerability toward the ROS-inducing drug Elesclomol.

Collectively, our results exemplify how aberrant activation of a developmental transcription factor by a dominant oncogene can promote malignancy, but provide opportunities for targeted therapy.

---

Ewing sarcoma (EwS) is the second most common bone or soft-tissue cancer in children and adolescents^1^. Even though the cell of origin of EwS is still debated, increasing evidence suggests that it may arise from mesenchymal stem cell (MSC)-derived early committed osteochondrogenic progenitors^2,3^. Indeed, EwS cells display a highly undifferentiated and embryonal phenotype. Clinically, EwS is a rapidly metastasizing cancer, and ~25% of cases are metastatic at initial diagnosis^1^. While great advances in treatment of localized disease have been achieved, established therapies still have limited success in advanced stages despite high toxicity^1^. Thus, more specific and in particular less toxic therapies are urgently required.

As a genetic hallmark, EwS tumors express chimeric EWSR1-ETS (EwS breakpoint region 1 – E26 transformation specific) fusion oncoproteins generated through fusion of the *EWSR1* gene and variable members of the ETS-family of transcription factors (TF), most commonly *FLI1* (85% of all cases)^4,5^. Prior studies demonstrated that *EWSR1-FLI1* acts as a pioneer transcription factor that massively rewires the tumor transcriptome ultimately promoting the malignant phenotype of EwS^6,7^. This is in part mediated through interference with and/or aberrant activation of developmental pathways^3,8^. Remarkably, EWSR1-FLI1 regulates approximately 40% of its target genes by binding to otherwise non-functional GGAA-microsatellites (mSats)^9^that are thereby converted into potent *de novo* enhancers, whose activity increases with the number of consecutive GGAA-repeats^7,10–14^.

Although EWSR1-FLI1 would in principle constitute a highly specific target for therapy, this fusion oncoprotein proved to be notoriously difficult to drug due to its intranuclear localization, its activity as a transcription factor^15,16^, the absence of regulatory protein residues^1^, its low immunogenicity^17^, and the high and ubiquitous expression of its constituting genes in adult tissues^1^. Hence, we reasoned that developmental genes and pathways that are aberrantly activated by EWSR1-FLI1 and virtually inactive in normal adult tissues, could constitute druggable surrogate targets.

As EwS most commonly arise in bone and possibly descend from osteo-chondrogenic progenitor cells,^3^ we speculated that EWSR1-FLI1 might interfere with bone developmental pathways. The transcription and splicing factor SOX6 (SRY-box 6) plays an important role in endochondral ossification^18^. Interestingly, its transient high expression delineates cells along the osteo-chondrogenic lineage showing high rates of proliferation while maintaining an immature phenotype along this lineage^19–22^.

In the current study, we show that EWSR1-FLI1 binds to an intronic GGAA-mSat within *SOX6*, which acts as an EWSR1-FLI1-dependent enhancer that induces the high and constitutive overexpression of SOX6 in EwS tumors. Moreover, we report that SOX6 promotes proliferation and tumorigenicity of EwS cells, and confers a druggable, therapeutic vulnerability toward the reactive oxygen species (ROS)-inducing small molecule Elesclomol, through upregulation of cell intrinsic ROS by interference with the antioxidant system.

## RESULTS

### SOX6 is highly but variably expressed in EwS

To explore the expression pattern of SOX6, we took advantage of a well-curated set of >750 DNA microarrays, which we established previously^23,24^, comprising 18 representative normal tissues types and 10 cancer entities. Comparative analyses revealed that SOX6 is overexpressed in EwS relative to normal tissues and other cancers (**Fig. 1a**). These data were validated on the protein level in a tissue microarray^23,24^ comprising the same normal tissue types and cancer entities (**Fig. 1b,c**). Both analyses showed that SOX6 is highly expressed in EwS tumors, albeit with substantial inter-tumor heterogeneity.

**Fig. 1.**
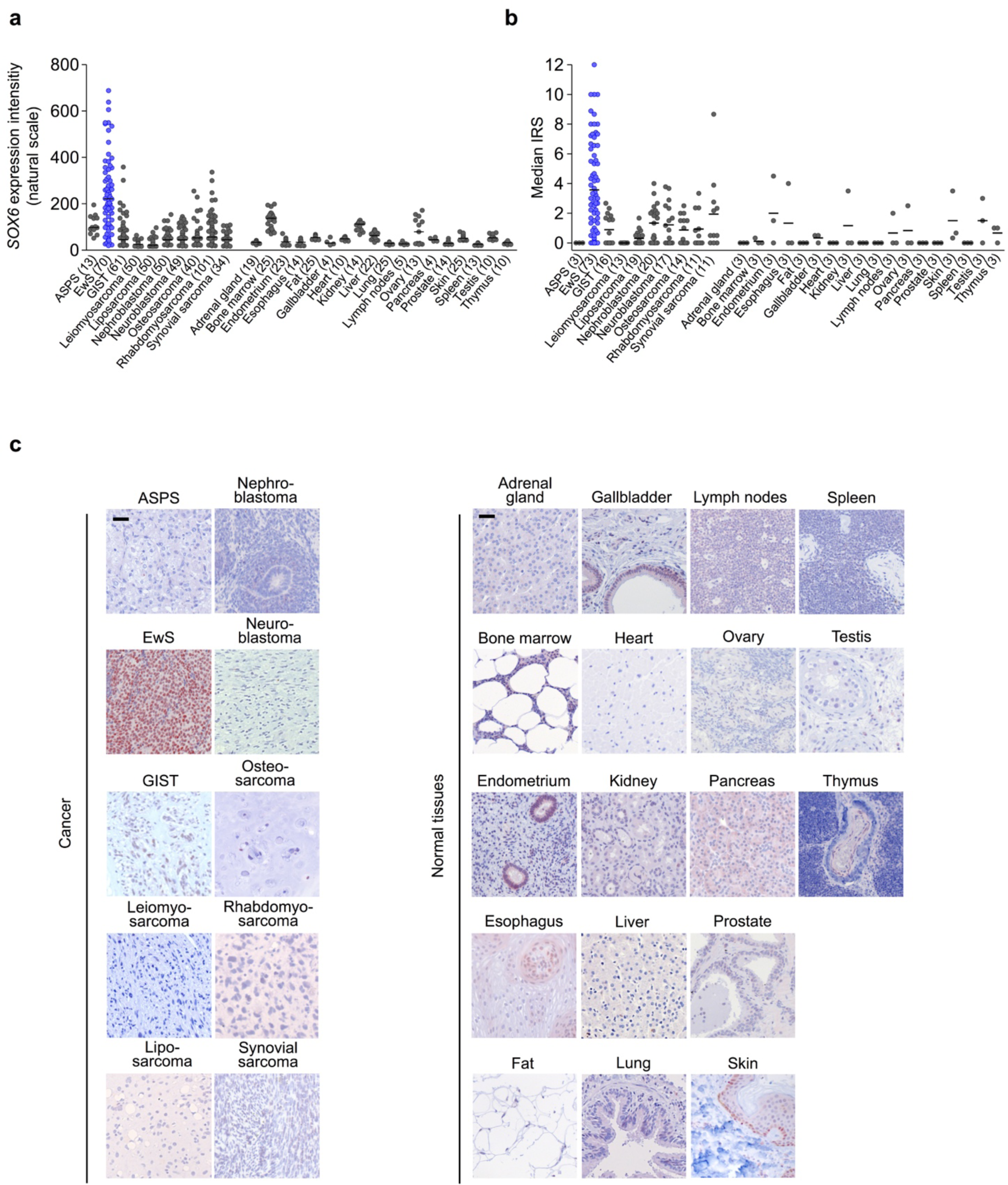
SOX6 is highly but variably expressed in EwS. **a)** Analysis of *SOX6* mRNA expression levels in EwS tumors, 9 additional sarcomas or pediatric tumors, and 18 normal tissue types as determined on Affymetrix HG-U133Plus2.0 arrays. Data are represented as dot plots and horizontal bars represent medians. The number of samples is given in parentheses. ASPS, alveolar soft part sarcoma; GIST, gastrointestinal stromal tumor. **b)** Validation of SOX6 expression on protein level by IHC in the same tissue types as shown in (**a**). Immuno Reactive scores (IRS) are presented as dot plots. Horizontal bars represent medians. The number of samples is given in parentheses. **c**) Representative micrographs of the IHC stains; scale bars=20 μm.

The generally high but variable expression of SOX6 was also observed in EwS cell line models compared to cell lines of three other pediatric cancer types including osteosarcoma (U2OS and SAOS-2), neuroblastoma (TGW and SK-N-AS) and rhabdomyosarcoma (Rh36 and Rh4) (**Supplementary Fig. 1**).

### *EWSR1-FLI1* induces *SOX6* expression via an intronic enhancer-like GGAA-mSat

The relatively high expression of *SOX6* in EwS compared to other sarcomas and pediatric cancers implied that there might be a regulatory relationship with the EwS specific fusion oncogene *EWSR1-FLI1*. Indeed, knockdown of *EWSR1-FLI1* in A673/TR/shEF1 and SK-N-MC/TR/shEF1 cells harboring a Dox-inducible short hairpin RNA (shRNA) against the fusion gene strongly reduced *SOX6* expression in a time-dependent manner *in vitro* (**Fig. 2a, b**, **Supplementary Fig. 2a**) and *in vivo* (**Fig. 2b**). Conversely, ectopic expression of *EWSR1-FLI1* in human embryoid bodies strongly induced *SOX6* expression (**Fig. 2c**).

**Fig. 2.**
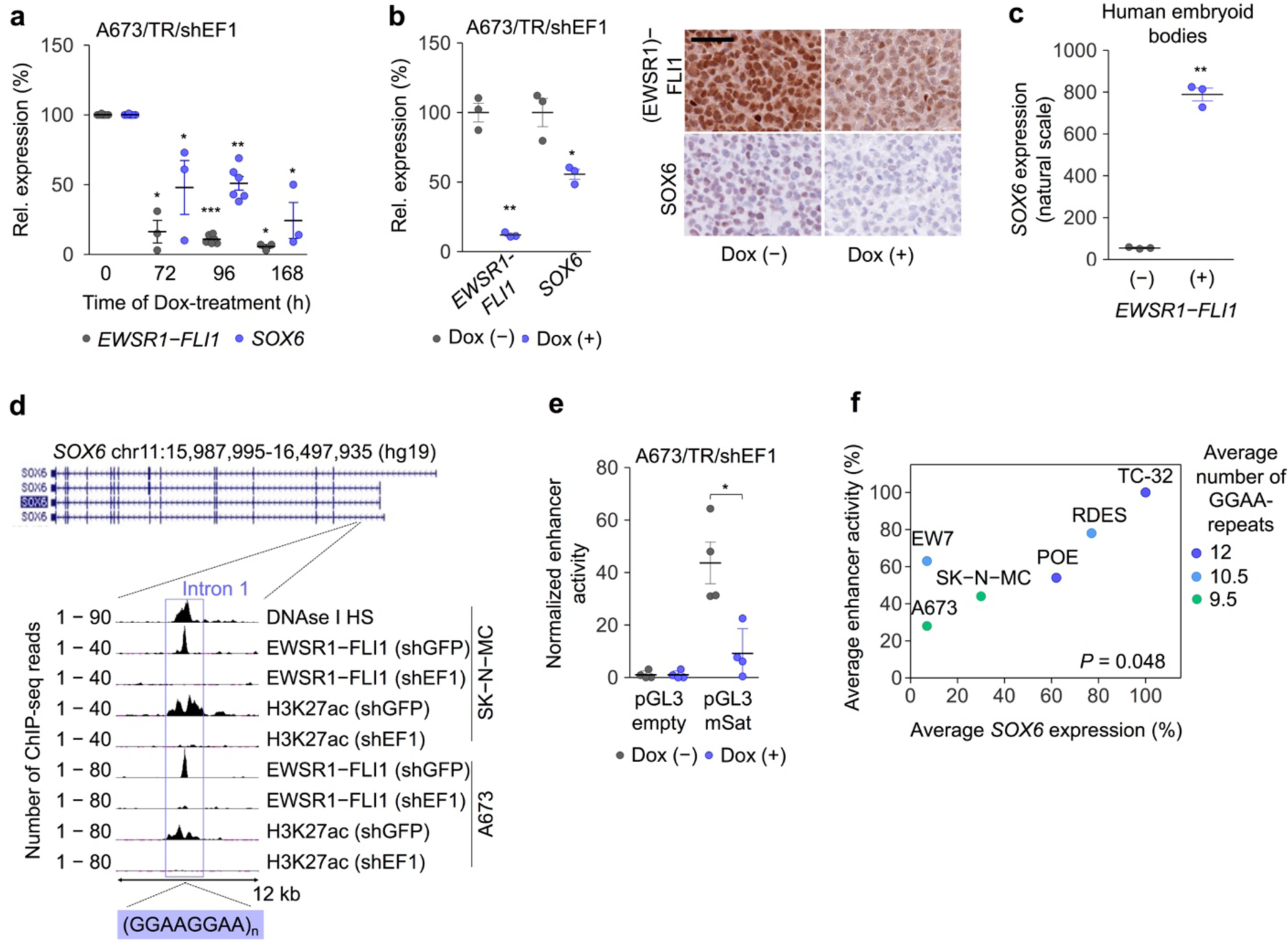
*EWSR1-FLI1* induces *SOX6* expression via an intronic enhancer-like GGAA-mSat. **a)** Analysis of *EWSR1-FLI1* and *SOX6* expression by qRT-PCR in A673/TR/shEF1 cells at indicated time points after addition of Dox. Horizontal bars represent means and whiskers SEM, *n*≥3. *P* values determined via two-sided Mann-Whitney test. **b)** Left: Analysis of *EWSR1-FLI1* and *SOX6* expression by Affymetrix microarrays in xenografts from A673/TR/shEF1 cells 96h after start of Dox-addition, Horizontal bars represent means and whiskers SEM, *n*=3. *P* value determined via independent one-sample t-test. Right: Representative immunohistological stains of xenografts stained for (EWSR1)FLI1 and SOX6. Scale bar=20µm. **c)** Analysis of *SOX6* expression by Affymetrix microarrays in embryoid bodies after ectopic *EWSR1-FLI1* expression. Horizontal bars represent means and whiskers SEM, *n*=3. *P* value determined via unpaired two-sided t-test with Welch’s correction. **d)** Integrated genomic view of the *SOX6* locus displaying tracks for DNAse 1 hypersensitivity (HS) and ChIP-Seq data for EWSR1-FLI1 and H3K27ac in A673 and SK-N-MC EwS cells transfected with shRNA against EWSR1-FLI1 (shEF1) or control shRNA (shGFP). **e)** Analysis of relative enhancer activity of the SOX6-associated GGAA-mSat by dual luciferase reporter assays in A673/TR/shEF1 cells (-/+). Horizontal bars represent means and whiskers SEM, *n*=4. *P* value determined via two-sided Mann-Whitney test. **f)** Correlation of the average enhancer activity of both alleles of the *SOX6-* associated GGAA-mSat and the average *SOX6* mRNA expression levels across six EwS cell lines (TC-32 was set as reference). The color code indicates the average number of consecutive GGAA-repeats of both alleles. \*\*\**P*<0.001, \*\**P*<0.01, \**P*<0.05

To investigate the underlying mechanism of this regulatory relationship, we analyzed publicly available DNase-Seq and ChIP-Seq data of two EwS cell lines (A673 and SK-N-MC) and found a prominent EWSR1-FLI1 peak within intron 1 of *SOX6*, which was strongly reduced upon *EWSR1-FLI1* knockdown (**Fig. 2d**). This EWSR1-FLI1 peak mapped to a GGAA-mSat located within a DNase 1 hypersensitivity site, indicating open chromatin, and showed EWSR1-FLI1-dependent acetylation of H3K27, which marks active enhancers (**Fig. 2d**). The EWSR1-FLI1-dependent enhancer activity of this GGAA-mSat was confirmed by luciferase reporter assays in A673/TR/shEF1 cells transfected with pGL3 reporter plasmids in which we cloned a 1-kb fragment containing this *SOX6-*associated GGAA-mSat from the human reference genome (**Fig. 2e**).

As prior studies showed that the enhancer activity at EWSR1-FLI1-bound GGAA-mSats positively correlates with the number of consecutive GGAA-repeats^12,25^, we hypothesized that the observed variability in *SOX6* expression might be caused by differences in repeat numbers at the *SOX6*-associated GGAA-mSat. To test this possibility, we cloned both alleles for this mSat from six EwS cell lines with largely different *SOX6* expression levels (**Supplementary Fig. 1**), determined their repeat number by Sanger sequencing, and measured their enhancer activity in reporter assays. We observed a positive correlation (*P* = 0.047) of the average *SOX6* expression levels with the observed average enhancer activity across cell lines, which corresponded to the average repeat numbers of both alleles (**Fig. 2f**, **Supplementary Table 1**). Interestingly, the inter-individual differences in *SOX6* expression levels correlated neither with (minor) differences of *SOX6* promoter methylation nor with copy number variations at the *SOX6* locus in primary EwS tumors (**Supplementary Fig. 2b, c**).

Collectively, these data suggest that *EWSR1-FLI1* induces *SOX6* by binding to a polymorphic intronic GGAA-mSat, which exhibits length-dependent enhancer activity.

### SOX6 promotes proliferation of EwS cells *in vitro* and *in vivo*

To explore the possible function of *SOX6* in EwS, we generated two cell lines (RDES and TC-32) with doxycycline (Dox)-inducible shRNAs against *SOX6* (shSOX6_2 and shSOX6_3) and corresponding controls with a Dox-inducible non-targeting control shRNA (shCtrl). In these transduced cells, addition of Dox (0.1 µg/ml) to the culture medium effectively silenced SOX6 expression at the mRNA and protein level (**Fig. 3a, Supplementary Fig. 3a**).

**Fig. 3.**
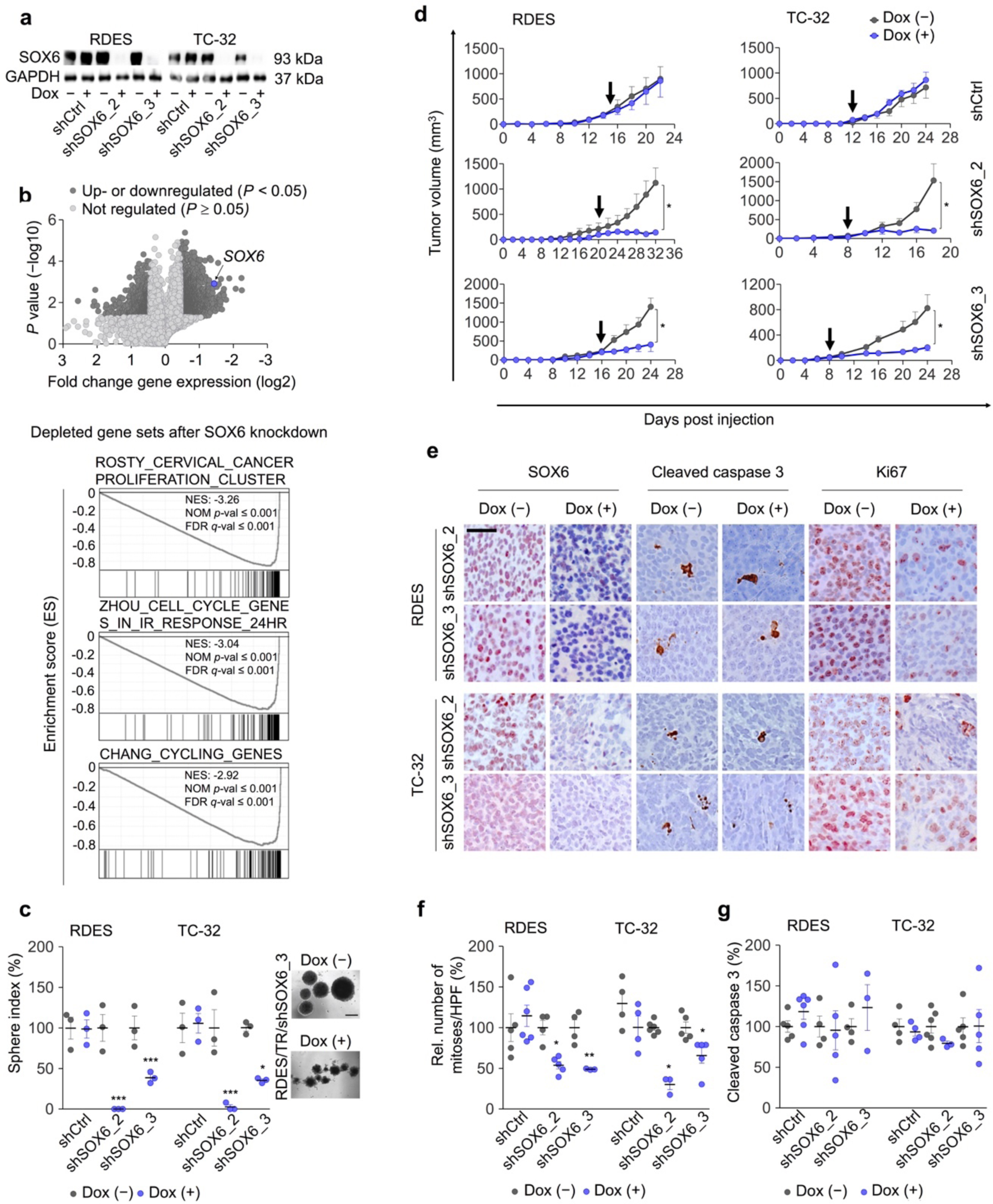
SOX6 promotes proliferation of EwS cells *in vitro* and *in vivo*. **a)** Western blot analysis 96h after Dox-induced shRNA-mediated SOX6 knockdown in RDES and TC-32 EwS cells. GAPDH served as loading control. **b)** Top: Volcano plot of microarray data showing differentially expressed genes (DEGs) after shRNA-mediated *SOX6* knockdown compared to a non-targeting shCtrl. A summary of two EwS cell lines is shown. Bottom: Representative enrichment plots from GSEA of transcriptome profiles of RDES and TC-32 EwS cells 96h after induction of shRNA-mediated SOX6 silencing. **c)** Left: Quantification of the sphere index after 12 days of Dox-treatment in RDES and TC-32 cells. Horizontal bars represent means and whiskers the SEM, *n*=3. *P* values determined via two-sided Mann-Whitney test. Right: Representative micrographs of RDES/TR/shSOX6_3 spheres. Scale bar=1 mm. **d)** Analysis of tumor growth of xenografted RDES and TC-32 cells containing either Dox-inducible specific shRNAs against *SOX6* (shSOX6_2/shSOX6_3) or a non-targeting control shRNA (shCtrl). When tumors were palpable (arrow), mice were randomized and henceforth treated with Dox (+) or vehicle (–). Data are represented as means and SEM, *n*≥3 mice per condition. *P* values determined via two-sided Mann-Whitney test. **e)** Representative micrographs of xenografts from (**d**) showing IHC stains for SOX6, cleaved caspase 3 and Ki67. Scale bar=20µm. **f)** Quantification of the relative number of mitoses per high-power field (HPF) of xenografts shown in (**d**). Horizontal bars represent means and whiskers the SEM, *n*≥3. *P* values determined via two-sided Mann-Whitney test. **g)** Quantification of the relative number of cells positive for cleaved caspase 3 of xenografts shown in (**d**). Horizontal bars represent means and whiskers the SEM, *n*≥3. \*\*\**P*<0.001, \*\**P*<0.01, \**P*<0.05.

Since SOX6 acts – depending on the cellular context – as a splicing and/or transcription factor^26,27^, we explored the effect of *SOX6* knockdown in RDES and TC-32 EwS cell lines using Affymetrix Clariom D arrays, which enable the simultaneous transcriptome-wide analysis of splicing events and differential gene expression. While the knockdown of *SOX6* for 96h had little effect on splicing (**Supplementary Table 2**), we noted a strong effect on differential gene expression (**Fig. 3b**). In fact, *SOX6* silencing induced a concordant up- or downregulation (FC<-0.5 and FC >+0.5; *P*<0.05) of 816 and 3,145 genes, respectively, across shRNAs and cell lines (**Supplementary Table 3**). Gene set enrichment analysis (GSEA) of these differentially expressed genes (DEGs) identified a strong depletion of proliferation-related gene signatures in SOX6 silenced EwS cells (**Fig. 3b, Supplementary Table 4**).

To validate the predicted role of SOX6 in EwS proliferation, we performed knockdown experiments using pooled short interfering RNAs (sipool) against *SOX6* in three EwS cell lines (POE, RDES, TC-32). Each sipool consisted of 30 different siRNAs, which virtually eliminates off-target effects^28^, and which induced a 60-80% *SOX6* knockdown as compared to a non-targeting control sipool (sipCtrl) after 96h. In these experiments, we noted a significant reduction of the viable cell count in all EwS cell lines in standardized cell counting experiments (including the supernatant) (**Supplementary Fig. 3b**). In accordance, Dox-induced long-term *SOX6* knockdown significantly reduced the 2D clonogenic and 3D sphere formation capacities of EwS cell lines as compared to controls (Dox (-) and shCtrl) (**Fig. 3c, Supplementary Fig. 3c**).

To test whether this effect was mediated via alteration of the cell cycle, we carried out flow cytometric assays with propidium iodine (PI). In serum-starved and thus G_0_-synchronized cells, we observed a significant delay in cell cycle progression 20h after re-addition of serum in *SOX6*-silenced cells (**Supplementary Fig. 3d**).

To assess the potential contribution of SOX6 to tumor growth of EwS cells *in vivo*, we performed xenograft experiments by injecting two different EwS cell lines with Dox-inducible shRNAs against *SOX6* subcutaneously into the flanks of NSG mice. While no effect of Dox-treatment was apparent in EwS cell lines expressing the non-targeting control shRNA, we noted a strong and consistent reduction of tumor growth upon *SOX6* knockdown in both shRNA constructs and both cell lines (**Fig. 3d**). The knockdown of *SOX6* was confirmed *ex vivo* in xenografts by qRT-PCR (**Fig. 3e**, **Supplementary Fig. 3e**) and by immunohistochemistry (IHC) (not shown). Immunohistological assessment showed that SOX6 silencing was associated with a significant reduction of proliferation as indicated by numbers of mitotic cells per high-power filed (HPF) and Ki67 stains (**Fig. 3f, Supplementary Fig. 3f**). In contrast, no significant differences in cleaved caspase 3 and Annexin V staining were observed (**Fig. 3g, Supplementary Fig. 3g**), suggesting that the apparent reduction of tumor growth was not mediated by apoptotic cell death.

Among the proliferation-associated genes downregulated after *SOX6* knockdown (**Fig. 3b**), three genes (*CDCA3, DEPDC1* and *E2F8*) appeared as plausible candidate genes to promote the pro-proliferative phenotype of SOX6, as they were previously shown to be involved in cell cycle progression^29–32^. In accordance, knockdown of any one of the genes with specific sipools in RDES and TC-32 EwS cells phenocopied, at least in part, the proliferative effect of SOX6 (**Supplementary Fig. 3h, i**).

Collectively, these results highlight a contribution of SOX6 to proliferation, clonogenic growth and tumorigenicity of EwS cells.

### High expression of SOX6 confers sensitivity toward the small-molecule Elesclomol in EwS

To explore whether high SOX6 levels could constitute a specific vulnerability of EwS that may be exploited therapeutically, we interrogated a published gene expression dataset with matched drug-response data comprising 22 EwS cell lines^33^. To this end, we calculated for all 264 tested drugs the Pearson correlation coefficient and its statistical significance of the corresponding IC50 values with the observed *SOX6* expression levels across EwS cell lines (**Fig. 4a**).

**Fig. 4.**
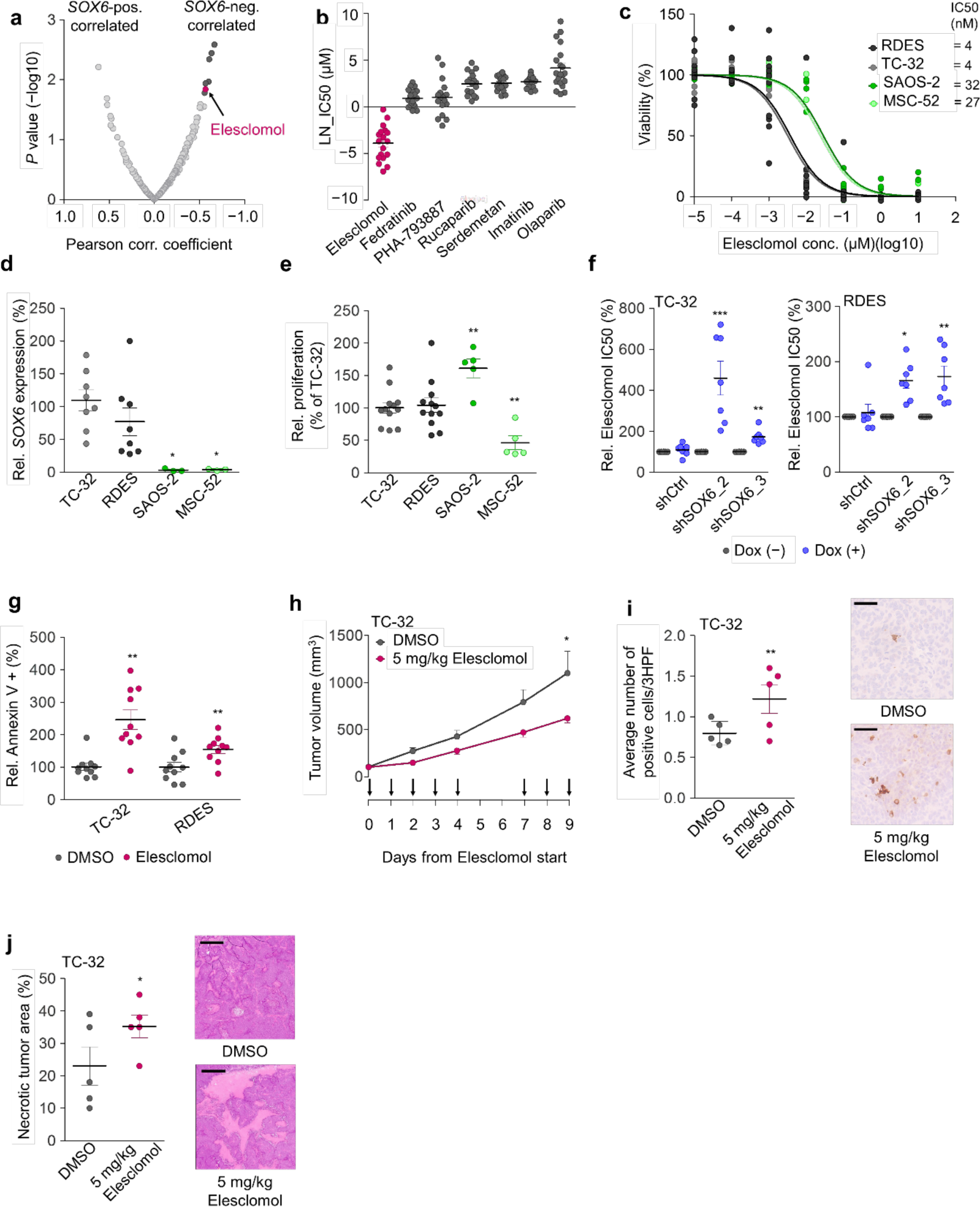
High expression of SOX6 confers sensitivity toward the small-molecule Elesclomol in EwS. **a)** Analysis of publicly available matched gene expression and drug-response data^33^ of up to 22 EwS cell lines per drug. Highlighted in dark grey, top 7 drugs with *P*<0.02; pink = Elesclomol. **b)** LN_IC50 (µM) of the top 7 drugs including Federatinib (JAK-2 inhibitor), PHA-793887 (CDK2/5/7 inhibitor), Rucaparib (PARP inhibitor), Serdemetan (p53 activator), Imatinib (tyrosine kinase inhibitor) and Olaparib (PARP1/2 inhibitor) with *P*<0.02. Horizontal bars represent means and whiskers SEM, *n*≥18 EwS cell lines. **c**) Quantification of relative viability of indicated cell lines by a Resazurin assay after treatment with Elesclomol at indicated concentrations for 72h. Modeled dose-response curves and calculated IC50 values (nM) are displayed for SOX6-high expressing EwS cells (black and grey) and the SOX6-low expressing osteosarcoma cell line SAOS-2 and the mesenchymal cell line MSC-52 (dark and light green), *n*≥3. **d**) Analysis of relative SOX6 expression in indicated cell lines by qRT-PCR. Horizontal bars represent means and whiskers SEM, *n*≥3. *P* values determined via two-sided Mann-Whitney test. **e**) Analysis of cell viability of indicated cell lines by a Resazurin assay. Horizontal bars represent means and whiskers SEM, *n*≥5. *P* values determined via two-sided Mann-Whitney test. **f**) Quantification of relative Elesclomol IC50 values by a Resazurin assay in indicated cell lines after 72h of Elesclomol treatment and concomitant addition of Dox. Horizontal bars represent means and whiskers SEM, *n*=7. *P* values determined via two-sided Mann-Whitney test. **g**) Quantification of relative Annexin V positivity of indicated EwS cells 48h after treatment with Elesclomol (10 nM). Horizontal bars represent means and whiskers SEM, *n*=10. *P* values determined via unpaired two-sided t-test with Welch’s correction. **h**) Analysis of tumor growth of TC-32 EwS cells in NSG mice treated once per day (day 0-4 and day 7-9) with Elesclomol (intravenously, 5 mg/kg). Data represent means and SEM, *n*=5 mice per condition. *P* values determined via two-sided Mann-Whitney test. **i**) Left: Quantification of the average number of cleaved caspase 3 positive cells per 3 HPF in TC-32 xenografts shown in (**h**). Horizontal bars represent means and whiskers SEM, *n*=5 per condition. *P* values determined via two-sided Mann-Whitney test. Right: representative micrographs. Scale bar=100 µm. **j**) Left: Quantification of necrotic area in TC-32 xenografts shown in (**h**). Horizontal bars represent means and whiskers SEM, *n*=5 per condition. *P* values determined via two-sided Mann-Whitney test. Right: Representative micrographs. Scale bar = 900 µm. \*\*\**P*<0.001, \*\**P*<0.01, \**P*<0.05.

Among the top 7 drugs, Elesclomol (N-malonyl-bis (N-methyl-N-thiobenzoyl hydrazide) (*r*_Pearson_= −0.565; *P*=0.014) was the only drug, which effectively could inhibit EwS growth at a nanomolar range (IC50 **~**20 nM) (**Fig. 4b**). Elesclomol is a potent oxidative stress inducer, which is believed to exert its pro-apoptotic effects in cancer cells via elevating ROS levels beyond a tolerable threshold^34^. Indeed, in validation drug-response assays, Elesclomol strongly decreased viability of EwS cells with high *SOX6* levels while the osteosarcoma cell line SAOS-2 and non-transformed human primary MSC line MSC-52 that exhibit low *SOX6* expression levels were relatively resistant (**Fig. 4c, d**). The high sensitivity of EwS cells toward Elesclomol appeared to be independent of proliferation under normal conditions, since the osteosarcoma cell line SAOS-2 proliferated even more than the tested EwS cells (**Fig. 4e**). Yet, knockdown of *SOX6* in RDES and TC-32 EwS cells significantly diminished their sensitivity toward Elesclomol (**Fig. 4f**), pointing to a functional role of SOX6 in Elesclomol-sensitivity. Consistent with a prior report in other cancer cell lines^34^, Elesclomol strongly induced apoptosis in EwS cell lines *in vitro* when treated with corresponding IC50 concentrations (**Fig. 4g**), without affecting *SOX6* expression levels (**Supplementary Fig. 4a**). In accordance, intravenous administration of Elesclomol for 9 days reduced local tumor growth of TC-32 EwS xenografts *in vivo* (**Fig. 4h**), which was accompanied by induction of apoptosis and cell death as evidenced by significantly increased numbers of cells positive for cleaved caspase 3 (**Fig. 4i**), and more necrotic tumor area (**Fig. 4j**). Of note, mice treated with Elesclomol did not exhibit overt adverse effects such as weight-loss (**Supplementary Fig. 4b**) or histo-morphological changes in the inner organs (not shown).

In sum, these results demonstrate that *SOX6* expression confers a proliferation-independent sensitivity toward the ROS-inducing small-molecule Elesclomol to EwS cells.

### SOX6 induces intracellular ROS through interference with the antioxidant system

Since Elesclomol can induce oxidative stress, we investigated whether Elesclomol treatment modulates ROS levels in EwS cell lines and why EwS cells are sensitive to Elesclomol. Indeed, treatment of RDES and TC-32 cells with Elesclomol (10 nM) significantly induced ROS in both EwS cell lines compared to control (DMSO) (**Fig. 5a**). To test whether ROS levels play a role in the capacity of Elesclomol to kill EwS cells, we carried out drug-response assays in the presence/absence of the antioxidant N-acetylcysteine (Nac), which is able to scavenge free radicals^35^. In both cell lines, Nac-treatment resulted in significantly increased IC50 values indicating that Elesclomol exerts its pro-apoptotic effect in EwS via ROS (**Fig. 5b**).

**Fig. 5.**
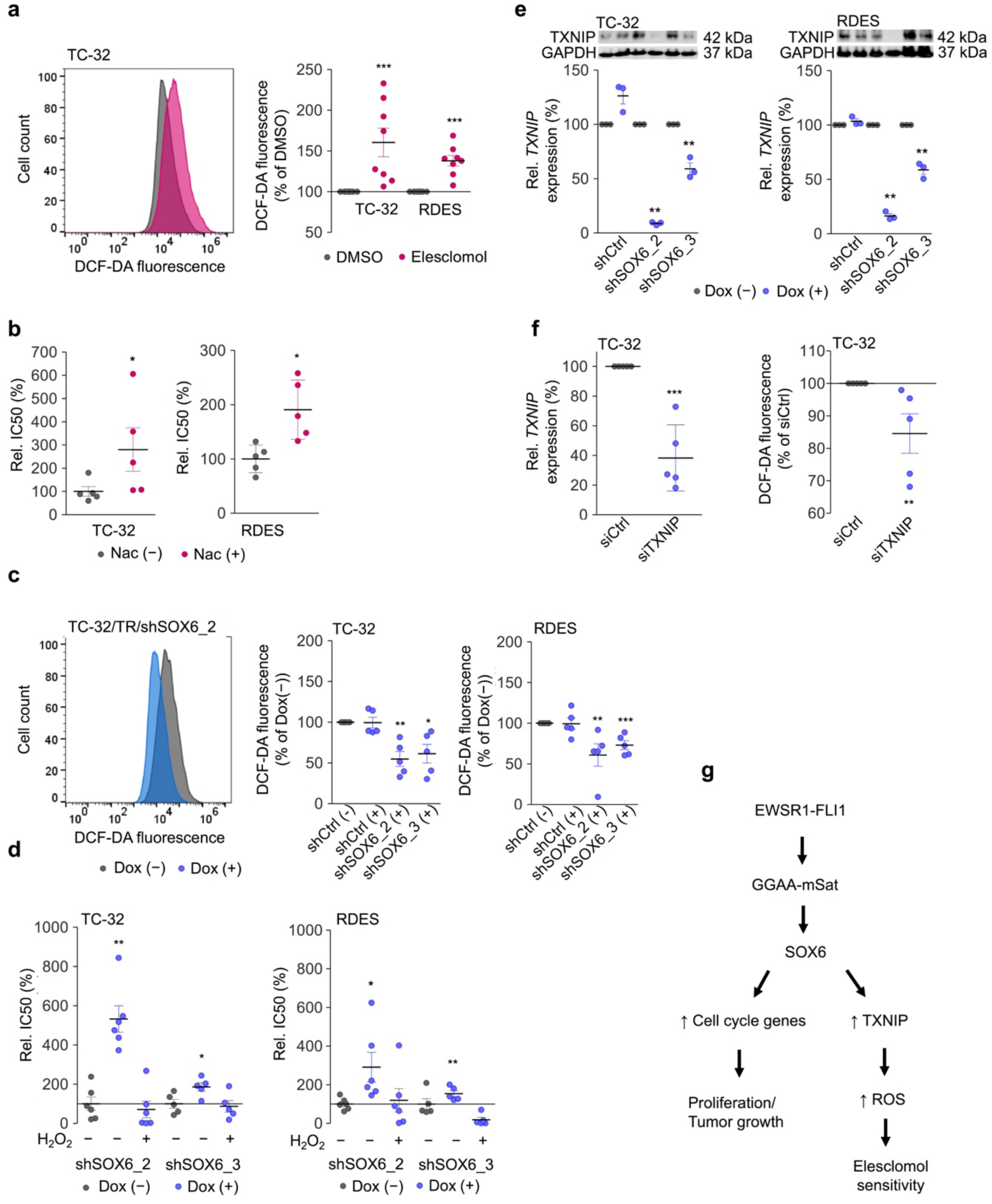
SOX6 induces intracellular ROS through interference with the antioxidant system. **a)** Left: Representative picture of flow cytometric measurement of ROS by DCF-DA fluorescence in TC-32 cells after Elesclomol-treatment (10 nM, pink color) compared to DMSO control (gray color). Right: Quantification of relative DCF-DA fluorescence in TC-32 and RDES cells after Elesclomol-treatment (10 nM, pink color) compared to DMSO control. Horizontal bars indicate means and whiskers SEM, *n*=8. *P* values determined via independent one sample t-test. **b)** Quantification of relative IC50 concentrations in TC-32 and RDES cells by a Resazurin assays after pre-treatment with the antioxidant N-acetylcysteine (Nac(+), pink color) compared to DMSO control (Nac(-), gray color). Horizontal bars indicate means and whiskers SEM, *n*≥5. *P* values determined via two-sided Mann-Whitney test. **c)** Left: Representative picture of flow cytometric measurement of ROS by DCF-DA fluorescence in TC-32/TR/shSOX6_2 cells after Dox-induced knockdown of SOX6 (blue color) for 96h as compared to control (Dox (–), gray color). Right: Quantification of relative DCF-DA fluorescence in TC-32 and RDES cells after SOX6 knockdown. Dox(–) = gray color, Dox(+) = blue color. Horizontal bars represent means and whiskers SEM, *n*≥3. *P* values determined via one-sample t-test. **d)** Quantification of relative IC50 Elesclomol concentrations in TC-32 and RDES cells by a Resazurin assays after Dox-induced SOX6 knockdown and treatment with H_2_O_2_ (30 µmol/l) for 72h. Horizontal bars represent means and whiskers SEM, *n*≥5. *P* values determined via two-sided Mann-Whitney test. **e)** Top: Representative Western blot analysis of TXNIP expression 96h after induction of SOX6 knockdown in TC-32 and RDES cells. GAPDH served as loading control. Bottom: Quantification of relative *TXNIP* mRNA expression in the same cells by qRT-PCR. Horizontal bars represent means and whiskers SEM, *n*=3. *P* values determined via two-sided Mann-Whitney test. **f)** Left: Analysis of relative *TXNIP* expression by qRT-PCR in TC-32 cells 96h after transfection with siRNA directed against *TXNIP*. Horizontal bars represent means and whiskers SEM, *n*=5. *P* value determined via unpaired two-sided t-test with Welch’s correction. Right: Analysis of ROS levels by flow cytometric measurement of DCF-DA fluorescence in TC-32 cells after siRNA-induced *TXNIP* knockdown. Horizontal bars represent means and whiskers SEM, *n*=5. *P* values determined via independent one sample t-test. **g)** Schematic illustration of the EWSR1-FLI1-mediated effect on SOX6 expression in EwS. \*\*\**P*<0.001, \*\**P*<0.01, \**P*<0.05

In line with this hypothesis, Dox-induced knockdown of SOX6 reduced ROS levels in both EwS cell lines (RDES and TC-32) and for both shRNA constructs, which was not observed in corresponding controls (shCtrl) (**Fig. 5c**). To functionally validate the ROS-dependent sensitivity toward Elesclomol conferred by SOX6 on EwS cells, we performed rescue experiments. In those we noted that addition of the potent ROS-inducer H_2_O_2_ on the SOX6 silenced EwS cells could fully restore the sensitivity of these cell lines toward Elesclomol (**Fig. 5d**), while having no effect on viability of EwS cells that were not treated with Elesclomol (**Supplementary Figure 5**).

These data suggest that SOX6 is involved in ROS metabolism and prompt further analysis of our available microarray data obtained from EwS cells with/without *SOX6* knockdown. Although we did not find evidence for a systematic enrichment/depletion of ROS-associated pathways in our GSEA, we identified *TXNIP* (thioredoxin interacting protein) – a key inhibitor of the thioredoxin antioxidant system – among the top 10 downregulated genes after *SOX6* silencing (**Supplementary Table 3**). The downregulation of TXNIP after silencing of SOX6 was confirmed in independent experiments on mRNA and protein level (**Fig. 5e**). Interestingly, knockdown of *TXNIP* in EwS cells reduced intracellular ROS levels (**Fig. 5f**).

Taken together, these data suggest that SOX6 interferes via TXNIP with the antioxidant system, which increases intracellular ROS levels and thus promotes Elesclomol sensitivity in EwS cells (**Fig. 5g**).

## DISCUSSION

EwS is a highly aggressive cancer, affecting bone or soft-tissue, possibly descending from chondro-/osteo-progenitors. Since the transcription factor SOX6 is crucial for endochondral ossification and thus for bone development^18,36^, we aimed at analyzing its role in EwS.

Our results show that *SOX6* is a direct EWSR1-FLI1 target gene that is highly but variably overexpressed on the mRNA and protein level in EwS as compared to most normal tissues and other cancers. However, we found no correlation of copy number variations and differences in promoter methylation with *SOX6* expression levels in EwS tumors. In contrast, we identified an intronic *SOX6*-associated GGAA-mSat that was bound by EWSR1-FLI1 *in vivo* and which exhibited strong length- and EWSR1-FLI1-dependent enhancer activity in EwS cell lines. Thus, it is tempting to speculate that the observed inter-tumor heterogeneity in SOX6 expression of EwS tumors and EwS cell lines is likely caused by inter-individual differences in the number of consecutive GGAA-repeats at this *SOX6*-associated GGAA-mSat. These findings are in line with recent observations for other EWSR1-FLI1 target genes such as *EGR2* and *NR0B1* whose variable expression in EwS tumors is caused by inter-individual differences in GGAA-repeat numbers of the corresponding enhancer-like GGAA-mSat^1,37^.

Depending on the cellular context, SOX6 may act as a splicing factor^26,27^ or as transcriptional regulator^36^. In transcriptome profiling experiments that comprised >285,000 transcripts and isoforms, we did not observe a strong contribution of SOX6 to alternative splicing in EwS.

Instead, we identified a broad deregulation of a large number of genes after SOX6 silencing, pointing to a more pronounced role of SOX6 as a transcription factor in EwS. Especially the downregulated DEGs after SOX6 knockdown were significantly enriched for gene sets involved in proliferation and cell cycle progression. These changes in the cellular transcriptome were mirrored in functional *in vitro* and *in vivo* experiments. The strong pro-tumorigenic function of SOX6 in EwS is intriguing in other cancer entities such as esophageal squamous cell carcinoma and hepatocellular carcinoma *SOX6* was reported to act as a tumor suppressor^38,39^. However, in EwS, knockdown of *SOX6* strongly reduced anchorage-independent growth and tumorigenicity, which was accompanied by delayed transition through cell cycle phases and reduced expression of the proliferation marker Ki67. Thus, our results suggest that SOX6 may also have oncogenic properties, and that its oncogenic or tumor-suppressive function may depend on the cellular context.

Since novel therapeutic options for EwS patients are urgently required^3^, we investigated whether the high expression of SOX6 in EwS may provide a vulnerability that could be exploited therapeutically. Indeed, we discovered that high expression of SOX6 confers hyper-sensitivity toward the small-molecule Elesclomol. While Elesclomol was shown to inhibit cancer cell growth *in vitro* at micro-molar concentrations in melanoma, breast cancer and leukemia cell lines^40–43^ we noted a much higher sensitivity of EwS cells toward Elesclomol with IC50 values in the nano-molar range. These observations suggest that the higher sensitivity of EwS toward Elesclomol may be caused by the relatively higher expression of *SOX6* in EwS compared to other cancers such as osteosarcoma and melanoma (**Supplementary Fig. 4c**). In support of this hypothesis, Elesclomol-treatment in combination with paclitaxel had only moderate effect on outcome of unselected patients affected by malignant melanoma in phase II and III clinical trials^44,45^, and may have shown higher efficacy when preselecting patients with higher SOX6 levels or higher intracellular ROS levels.

Since many cancer types including EwS display an oxidative stress phenotype characterized by higher ROS levels than normal tissues^46,47^, cancer cells tend to be more sensitive toward further increases in oxidative stress as nonmalignant cells^48^. Previous reports demonstrated that Elesclomol can induce ROS levels beyond a tolerable threshold triggering apoptosis^34,41,49^. In line with these findings, we observed an increase of ROS followed by apoptosis after Elesclomol-treatment *in vitro* and *in vivo* in EwS cells. The apoptotic effect appeared to be dependent on the baseline intracellular ROS levels since knockdown of SOX6 and concomitant downregulation of ROS or pre-treatment of EwS cells with the antioxidant Nac diminished the sensitivity of EwS cells toward Elesclomol, whereas addition of the ROS-inducer H_2_O_2_ rescued the SOX6 knockdown effect on Elesclomol sensitivity.

The elevated intracellular ROS levels and associated hyper-sensitivity of SOX6 high expressing EwS cells toward Elesclomol can be explained, at least in part, by the SOX6-mediated upregulation of TXNIP, an inhibitor of the thioredoxin (TRX) antioxidant system that plays an essential role in buffering intracellular ROS levels^50,51^.

These data suggest that ROS-inducing drugs such as Elesclomol could offer a new therapeutic option for EwS patients with high SOX6 expression levels. Additionally, SOX6 may serve as a biomarker to predict the efficacy of Elesclomol treatment in EwS, and perhaps other cancer types. Interestingly, Elesclomol treatment has been shown to potentiate the pro-apoptotic effect of ROS-dependent chemotherapeutic drugs such as doxorubicin in breast cancer^42^. Since doxorubicin is part of current standard treatment regimens for EwS patients, it is tempting to speculate that Elesclomol treatment may also serve as an enhancer for doxorubicin treatment, even in patients with relatively low intratumoral SOX6 levels.

In synopsis, we discovered that EWSR1-FLI1 hijacks SOX6 in EwS, which promotes tumor growth, and interferes with the antioxidant system creating a therapeutic vulnerability toward ROS-inducing drugs. Our results exemplify how aberrant activation of a developmental transcription factor by a dominant oncogene can promote malignancy but provide opportunities for targeted therapy.

## MATERIAL AND METHODS

### Cell lines and cell culture conditions

The neuroblastoma cell line SK-N-AS as well as HEK293T were purchased from ATCC. Human EwS cell lines and other cell lines were provided by the following repositories and/or sources: A673 and cells were purchased from ATCC. MHH-ES1, RDES, SK-N-MC and cells were provided by the German collection of Microorganism and Cell Cultures (DSMZ). CHLA-10, CHLA-25, CHLA-32, CHLA-57, CHLA-99, COG E-352, TC-32, TC-71 and TC-106 cells were kindly provided by the Children’s Oncology Group (COG), and ES7, EW1, EW3, EW7, EW16, EW17, EW18, EW22, EW24, ORS, POE, SK-PN-DW, SK-PN-LI and STA-ET1 as well as neuroblastoma (Rh4, Rh36) and rhabdomyosarcoma (TGW) cell lines cells were provided by O. Delattre (Institute Curie, Paris). A673/TR/shEF1 cells were provided by J. Alonso (Madrid, Spain)^21^. Human osteosarcoma cell lines SAOS-2 and U2OS were provided by DMSZ, and the MSC cell line MSC-52 was generated from bone marrow of a EwS patient (provided by U. Dirksen; Essen, Germany). All cell lines were cultured in RPMI-1640 medium with stable glutamine (Biochrom, Germany) supplemented with 10% tetracycline-free fetal bovine serum (Sigma-Aldrich, Germany), 100 U/ml penicillin and 100 µg/ml streptomycin (Merck, Germany) at 37°C with 5% CO_2_ in a humidified atmosphere. Cell lines were routinely tested for mycoplasma contamination by nested PCR, and cell line identity was regularly verified by STR-profiling.

### RNA extraction, reverse transcription, and quantitative real-time polymerase chain reaction (qRT-PCR)

Total RNA was isolated using the NucleoSpin RNA kit (Macherey-Nagel, Germany). 1 µg of total RNA was reverse-transcribed using High-Capacity cDNA Reverse Transcription Kit (Applied Biosystems, USA). qRT-PCR reactions were performed using SYBR green Mastermix (Applied Biosystems) mixed with diluted cDNA (1:10) and 0.5 µM forward and reverse primer on a BioRad CFX Connect instrument and analyzed using BioRad CFX Manager 3.1 software. Gene expression values were calculated using the 2^-(ΔΔCt) method^52^ relative to the housekeeping gene *RPLP0* as internal control. Oligonucleotides were purchased from MWG Eurofins Genomics (Germany) and are listed in **Supplementary Table 5**. The thermal conditions for qRT-PCR were as follows: heat activation at 95°C for 2 min, DNA denaturation at 95°C for 10 sec, and annealing and elongation at 60°C for 20 sec (50 cycles), final denaturation at 95°C for 30 sec.

### Transient RNA interference (RNAi)

POE, RDES and TC-32 cells were transiently reversely transfected with Lipofectamine RNAiMAX according to the manufacturer’s protocol (Invitrogen, USA) with either a non-targeting control sipool (sipCtrl) or sipools specifically directed against *CDCA3, DEPDC1, E2F8* or *SOX6* (all siTOOLs, Biotech, Germany) or short interfering RNA (siRNA) against *TXNIP* (MWG Eurofins, Germany) (**Supplementary Table 5**) at a final concentration of 5–15 nM depending on the cell line and target gene. Cells were re-transfected 48h after the first transfection. All sipools consisted of 30 different siRNAs directed against the target transcript, which eliminates off-target effects^28^. Knockdown efficacy was validated by qRT-PCR and Western blot.

### Cloning of GGAA-mSats and luciferase reporter assays

For luciferase reporter assays, both alleles of a 1 kb fragment including the *SOX6-*associated GGAA-mSat (hg19 coordinates: chr11: 16,394,850-16,395,749) were cloned from three highly *SOX6* expressing EwS cell lines (RDES, TC-32, POE) and three *SOX6* moderately/lowly expressing EwS cells (A673, SK-N-MC, EW7) using GoTaq DNA Polymerase (Promega, Germany) and primer sequences listed in **Supplementary Table 5**. The thermal conditions for touchdown (TD)-PCR protocol were as follows: Initial denaturation: 95°C for 2 min; denaturation 98°C for 10 sec; annealing: 59°-49°C for 30 sec (T_m_ (= 59°C) −5°C + 10°C higher for TD-PCR); −1°C every 2 cycles); extension: 72°C for 1 min (2х10 cycles, and subsequently another 20 cycles at an annealing temperature of 65°C); final extension: 72°C for 5 min. These fragments were cloned upstream of the SV40 minimal promoter into the pGL3 luciferase reporter vector (Promega, #E1761, Germany) by In-Fusion HD Cloning Kit (Clontech, Japan) according to the manufacturer’s protocol. These vectors were used for transformation of Stellar Competent Cells (Clontech); successfully transformed bacteria were selected using ampicillin (Merck). Correct insertion of the vector was confirmed by colony PCR. The number of consecutive GGAA-repeats for each allele and each cell line was determined by commercial Sanger sequencing using the following primer: 5′-CTTTATGTTTTTGGCGTCTTCCA-3’. For reporter assays, 5×10^4^ A673/TR/shEF1 cells per 24-well were seeded in 600 µl medium and transfected with pGL3 luciferase reporter vector and Renilla pGL3-Rluc (ratio, 100:1) using Lipofectamine LTX Reagent with PLUS Reagent (Invitrogen). Four hours after transfection, the culture media was replaced by media with/without Doxycycline (Dox; 1 µg/ml; Merck). Cells were lysed after 72h and monitored with a dual luciferase assay system (Berthold, Germany). Firefly luciferase activity was normalized to Renilla luciferase activity.

### Analysis of copy-number-variation (CNV) and promoter methylation in primary EwS

For CNV analysis, publicly available DNA copy number data for EwS tumors^53^ with corresponding RNA expression data (GSE34620 and GSE37371, *n*=32), were downloaded from the ‘soft tissue cancer – Ewing sarcoma – FR’ project from the International Cancer Genome Consortium (ICGC) Data Portal and Gene Expression Omnibus (GEO) of the NCBI, respectively. For the *SOX6* locus, segment mean values were extracted from these data using Visual Basic for Applications (VBA). The segment mean values were correlated with the log2-transformed expression of the candidate gene. For CpG methylation analysis, publicly available data on CpG methylation in 40 EwS tumors (GSE88826)^54^ with corresponding RNA expression data (GSE34620) were downloaded from GEO. For the *SOX6* locus, the ratio of methylated versus unmethylated reads was calculated for two CpG sites (CpG1 Hg19: chr11:15994482; CpG2 Hg19: chr11:15994519) in each sample (*n*=40) using VBA, which were covered by at least four reads.

### Analysis of *SOX6* expression levels in human embryoid bodies

Publicly available gene expression microarray data for ectopic EWSR1-FLI1 expression in human embryoid bodies generated on the Affymetrix HG-U133Plus2.0 array (GSE64686)^55^ were normalized by Robust Multiarray Average (RMA)^56^ using custom brainarray chip description files (CDF; ENTREZG, v19) yielding one optimized probe-set per gene^57^.

### Analysis of published DNase sequencing (DNase-Seq) and chromatin immune-precipitation followed by high-throughput DNA sequencing (ChIP-Seq) data

ENCODE SK-N-MC DNase-Seq (GSM736570) and ChIP-Seq data (GSE61944) were downloaded from the GEO, processed as previously described^25^ and displayed in the UCSC genome browser. The following samples were used in this study:

ENCODE_SKNMC_hg19_DNAseHS_rep1

GSM1517546_SKNMC.shGFP96.FLI1

GSM1517555_SKNMC.shFLI196.FLI1

GSM1517547_SKNMC.shGFP96.H3K27ac

GSM1517556_SKNMC.shFLI196.H3K27ac

GSM1517569_A673.shGFP48.FLI1

GSM1517572_A673.shFLI148.FLI1

GSM1517570_A673.shGFP48.H3K27ac

GSM1517573_A673.shFLI148.H3K27ac

### Transcriptome and splicing analyses

To asses an impact of SOX6 on gene expression and on alternative splicing in EwS, microarray analysis was performed. To this end, 1.2×10^4^ cells were seeded in wells of 6-well plates and treated with Dox for 96h (Dox-refreshment after 48h). Thereafter, total RNA was extracted with the ReliaPrep RNA Cell Miniprep System (Promega) and transcriptome profiled at IMGM laboratories (Germany). RNA quality was assessed with a Bioanalyzer and samples with RNA integrity numbers (RIN)>9 were hybridized to Human Affymetrix Clariom D microarrays. Data were log2-transformed quantile normalized with Affymetrix Expression Console Software (v1.4) using the SST-RMA algorithm as previously described^58^. Annotation of the data was performed using the Affymetrix library for Clariom D Array (human), both on gene and exon level. DEGs with consistent and significant fold changes (FCs) across shRNAs and cell lines were identified as follows: First, the FCs of the shControl samples and both specific shRNAs were calculated for each cell line separately. Then the FCs observed in the respective shControl samples were subtracted from those seen in the shSOX6 samples, which yielded the final FCs for both specific shRNAs. Then these final FCs for both specific shRNAs were averaged across cell lines to obtain the mean final FC per gene across shRNAs and cell lines. The consistency of differential gene expression was determined by an independent one-sample t-test. Only those genes were considered a DEG, which showed a minimum log2 FC>±0.5 and *P*<0.05. To identify enriched gene sets, genes were ranked by their expression FC between the groups Dox (-/+), and a pre-ranked GSEA (MSigDB v5.2, c2.cpg.all) with 1,000 permutations was performed^59^.

To assess the potential role of SOX6 in alternative splicing, we assumed that, in case RNA is not alternatively spliced, the ratio of each probe selection region (PSR) expression measured on the Affymetrix microarray with and without SOX6 knockdown stays the same independently of up- or downregulation of the corresponding gene. Hence, we calculated the additional FC between the expression value of each PSR before and after SOX6 knockdown to the FC expected by expression regulation assessed on the gene level. Of 539,385 PSRs with 47,851 matched genes in our analysis, 22,155 PSRs (10,754 genes) showed a consistent positive or negative additional log2 transformed expression FC of ≥0.3. For 20,050 PSRs (10,179 genes) expression differences were significant (*P*<0.05) when corrected for the FC on gene level. However, none of the PSRs remained significant after Bonferroni correction for multiple testing. Gene expression data were deposited at the GEO (accession code GSE120576).

### Generation of Doxycycline (Dox)-inducible short hairpin RNA (shRNA) expressing cells

For long-term experiments human EwS cell lines SK-N-MC, RDES and TC-32 were transduced with lentiviral pLKO-TET-ON all-in-one vector system (Plasmid #21915, Addgene) containing a puromycin resistance cassette, and a tet-responsive element for Dox-inducible expression of shRNA against *EWSR1-FLI1* (shEF1), *SOX6* (shSOX6) or a non-targeting control shRNA (shCtrl). Dox-inducible vectors were generated according to a publicly available protocol^60^ using In-Fusion HD Cloning Kit (Clontech) (**Supplementary Table 5**). Vectors were amplified in Stellar Competent Cells (Clontech) and integrated shRNA was verified by Sanger sequencing (primer: 5’-GGCAGGGATATTCACCATTATCGTTTCAGA-3’). Lentiviral particles were generated in HEK293T cells. Virus-containing supernatant was collected to infect the human EwS cell lines. Successfully infected cells were selected with 1.5 µg/ml puromycin (InVivoGen, USA). The shRNA expression for *SOX6* knockdown in EwS cells was achieved by adding 0.1 µg/ml Dox every 48h to the medium. Generated cell lines were designated as SK-N-MC/TR/shEF1, RDES/TR/shCtrl, RDES/TR/shSOX6_2, RDES/TR/shSOX6_3, TC-32/TR/shCtrl, TC-32/TR/shSOX6_2, and TC-32/TR/shSOX6_3.

### Western blot

RDES/TR/shCtrl, RDES/TR/shSOX6_2, RDES/TR/shSOX6_3, TC-32/TR/shCtrl, TC-32/TR/shSOX6_2, and TC-32/TR/shSOX6_3 EwS cells were treated for 96h with Dox to induce SOX6 knockdown. Whole cellular protein was extracted with RIPA buffer containing 1 mM Na_3_VO_4_ (Sigma-Aldrich) and protease inhibitor cocktail (Sigma-Aldrich). Western blots were incubated with mouse monoclonal anti-SOX6 antibody (1:1,000, sc-393314, Santa Cruz, Germany), rabbit monoclonal anti-TXNIP antibody (1:1,000, ab188865, Abcam, UK) and mouse monoclonal anti-GAPDH (1:800, sc-32233, Santa Cruz). The nitrocellulose membranes (GE Healthcare BioSciences, Germany) were secondary incubated with anti-mouse IgG (H+L) horseradish peroxidase coupled (1:3,000, W402b, Promega) and polyclonal anti-rabbit IgG (1:5000, R1364HRP, OriGene, Germany). Proteins were detected using chemiluminescence HRP substrate (Merck). Densitometric protein quantifications were carried out by ImageJ.

### Proliferation assays

For proliferation assays, 2×10^5^ EwS cells were seeded in wells of 6-well plates and treated with 0.1 µg/ml Dox every 48h for knockdown or transiently transfected with Lipofectamine RNAiMAX (Invitrogen, USA) and the respective sipool every 48h for a total period of 96h. Cell viability was determined including the supernatant by counting the cells with Trypan-Blue (Sigma-Aldrich) in standardized hemocytometers (C-Chip, Biochrom).

### Clonogenic growth assays

For clonogenic growth assays, RDES and TC-32 harboring shRNAs against *SOX6* were seeded at low density (200 cells) per well of a 12-well plate and grown for 21 days with renewal of Dox every 48h. The colonies were counted in three technical replicates and the colony area was measured with the ImageJ Plugin *Colony area*. The clonogenicity index was calculated by multiplying the counted colonies with the corresponding colony area.

### Sphere formation assay

For the analysis of anchorage-independent growth, EwS cell lines RDES and TC-32 harboring Dox-inducible shRNAs against *SOX6*, were pre-treated with Dox for 48h before seeding. Then, 1×10^3^ cells/96-well were seeded in Costar Ultra-low attachment plates (Corning, Germany) for 12 days. 20 µl of fresh medium with/without Dox was added every 48h. At day 12, wells were photographed and spheres larger than 500 µm in diameter were counted. The area was measured using ImageJ. The sphere volumes were calculated as follows: V = 4/3×π×r^3^. The sphere index was calculated by multiplying the counted colonies with the corresponding colony volume.

### Cell cycle analysis

For cell cycle analysis, RDES and TC-32 cells harboring a Dox-inducible shRNA against *SOX6* were seeded at 4×10^5^ cells per 10 cm dish and subsequently starved for 56h. Stimulation of the cells was performed with 10% FCS for 20h. Cells were fixed with ice-cold 70% ethanol, treated with 100 µg/ml RNAse (ThermoFisher, USA) and stained with 50 µg/ml propidium iodide (Sigma Aldrich). Analysis of the cell cycle was performed with BD Accuri C6 Cytometer (BD Biosciences) by counting at least 1×10^5^ events. An example for the gating strategy is provided in **Supplementary Figure 6a**.

### Annexin V staining

For analysis of Annexin V positive cells, RDES and TC-32 cells harboring a Dox-inducible shRNA against *SOX6* were seeded at 3×10^5^ cells/10 cm dish and treated with 0.1 µg/ml Dox every 48h for knockdown. After 96h, cells were washed with PBS and cells were resuspended in 1xAnnexin V buffer (BD Biosciences) with 5µl of Annexin V and 5µl PI solution for further 15 minutes. Analysis of Annexin V positivity was performed with BD Accuri C6 Cytometer (BD Biosciences) by counting at least 1×10^5^ events. An example for the gating strategy is provided in **Supplementary Figure 6b**.

### Reactive oxygen species (ROS) detection via DCF-DA fluorescence

For detection of ROS changes after *SOX6* knockdown, EwS cells were seeded at a density of 5×10^4^ cells/2 ml per 6-well and directly treated for 96h with Dox to induce the knockdown. For the knockdown of *TXNIP*, TC-32 wild type cells were seeded at a density of 7×10^4^ cells/2 ml per 6-well and reversely transfected with siRNA against *TXNIP* for 72h. At the day of analysis, cells were incubated in their medium with 2.5 µM DCF-DA (ThermoFisher) for 30 min at 37°C. Afterwards, cells were harvested and resuspended in PBS for flow cytometry analysis with Accuri C6 Cytometer (BD Biosciences). Gating strategy is provided in **Supplementary Figure 6c**.

### Gene expression and drug response correlation

To identify drugs whose efficacy correlates with *SOX6* expression in EwS cells, publicly available EwS cell line gene expression microarray data and drug-response values were downloaded from the EBI (E-MTAB-3610) and from www.cancerrxgene.org^33^. All CEL-files generated on Affymetrix Human Genome U219 arrays were simultaneously normalized using RMA^56^ and a custom brainarray chip description file (v20, ENTREZG) yielding one optimized probe set for each gene^57^. For all drugs tested in EwS cell lines, the Pearson correlation coefficient and its significance between *SOX6* expression and LN_IC50 values were calculated. Besides a high negative correlation coefficient and significance level, low IC50 values were chosen as criteria for selection of plausible and potentially relevant gene expression-drug response dependencies.

### Drug-response assays and Elesclomol treatment

For Elesclomol treatment, 2.5×10^3^ cells of RDES and TC-32 with Dox-inducible SOX6 knockdown as well as MSC-52 and SAOS-2 cell lines were seeded in wells of 96-well plates. Cells were pre-treated for 48h with Dox to induce *SOX6* knockdown before addition of Elesclomol (STA-4783) (Selleckchem, Germany). Different concentrations of Elesclomol ranging from 0.1 nM to 10 µM with/without Dox were added in a total volume of 100 µl per technical replicate for further 72h. For ROS scavenging experiments with N-acetylcysteine (Nac) (Sigma-Aldrich), cells were additionally treated with 0.01 mM Nac for 72h. For the rescue experiments with H_2_O_2_, 2.5×10^3^ cells/well were seeded in 96-well plates and *SOX6* knockdown was induced by addition of Dox. After 24h, cells were either treated with Elesclomol (10 nM) or vehicle, and in addition with 30 mmol/l H_2_O_2_. At the day of evaluation, Resazurin (16 µg/ml; Sigma-Aldrich) was added in order to measure cell viability. The relative IC50 concentrations were calculated using PRISM 5 (GraphPad Software Inc., CA, USA) and normalized to the respective controls.

### *In vivo* experiments

3×10^6^ EwS cells harboring a shRNA against *SOX6* were injected in a 1:1 mix of cells suspended in PBS with Geltrex Basement Membrane Mix (ThermoFisher) in the right flank of 10–12 weeks old NOD/Scid/gamma (NSG) mice. Tumor diameters were measured every second day with a caliper and tumor volume was calculated by the formula L×l^2^/2. When the tumors reached an average volume of 80 mm^3^, mice were randomized in two groups of which one henceforth was treated with 2 mg/ml BelaDox (Bela-pharm, Germany) dissolved in drinking water containing 5% sucrose (Sigma-Aldrich) to induce an *in vivo* knockdown (Dox (+)), whereas the other group only received 5% sucrose (control, Dox (-)). Once tumors of control groups reached an average volume of 1,500 mm^3^, all mice of the experiment were sacrificed by cervical dislocation. Other humane endpoints were determined as follows: Ulcerated tumors, loss of 20% body weight, constant curved or crouched body posture, bloody diarrhea or rectal prolapse, abnormal breathing, severe dehydration, visible abdominal distention, obese Body Condition Scores (BCS), apathy, and self-isolation. For *in vivo* experiments using Elesclomol, EwS cells were subcutaneously injected in mice as described above. When the tumors reached an average volume of 80 mm^3^, mice were randomly distributed in equal groups and henceforth treated once per day intravenously (i.v.) with 5 mg/kg Elesclomol or vehicle (DMSO), interrupted for a two-day break on days 6 and 7 to allow mice to recover from the i.v. injections. All tumor-bearing mice were sacrificed by cervical dislocation at the predefined experimental end-point, when 40% of control tumors exceeded a volume of 1,500 mm^3^. The tumors were extracted, small piece was snap frozen in liquid nitrogen for RNA isolation and the remaining tumor tissue was fixed in 4%-formalin and paraffin-embedded for immunohistology. Animal experiments were approved by the government of Upper Bavaria and conducted in accordance with ARRIVE guidelines, recommendations of the European Community (86/609/EEC), and UKCCCR (guidelines for the welfare and use of animals in cancer research).

### Human samples and ethics approval

Human tissue samples were retrieved from the archives of the Institute of Pathology of the LMU Munich (Germany) with approval of the institutional review board. The ethics committee of the LMU Munich approved the current study (approval no. 18-481 UE).

### Immunohistochemistry (IHC) and immunoreactivity scoring (IRS)

For IHC, 4-µm sections were cut and antigen retrieval was carried out by heat treatment with Target Retrieval Solution (S1699, Agilent Technologies, Germany). The slides were stained with either polyclonal anti-SOX6 antibody raised in rabbit (1:1,600; HPA003908, Atlas Antibodies, Sweden) or with monoclonal anti-Ki67 raised in rabbit (1:200, 275R-15, Cell Marque/Sigma-Aldrich) for 60 min at RT, followed by a monoclonal secondary horseradish peroxidase (HRP)-coupled horse-anti-rabbit antibody (ImmPRESS Reagent Kit, MP-7401, Vector Laboratories, Germany). AEC-Plus (K3469, Agilent Technologies) was used as chromogen. Samples were counterstained with hematoxylin (H-3401, Vector Laboratories). For cleaved caspase 3 staining, antigen retrieval was carried out by heat treatment with Target Retrieval Solution Citrate pH6 (S2369, Agilent Technologies). Slides were incubated with the polyclonal cleaved caspase 3 primary antibody (rabbit, 1:100; 9661, Cell Signaling, Frankfurt am Main, Germany) for 60 min at RT followed by ImmPRESS Reagent Kit. DAB+ (K3468, Agilent Technologies) was used as chromogen and hematoxylin for counterstaining. Formalin-fixed and paraffin-embedded (FFPE) xenografts of the EwS cell lines were stained with hematoxylin and eosin (H&E) for mitosis counting. Evaluation of immunoreactivity of SOX6 was carried out in analogy to scoring of hormone receptor Immune Reactive Score (IRS) ranging from 0–12 as previously described^23^. The percentage of cells with expression of the given antigen was scored and classified in five grades (grade 0 = 0-19%, grade 1 = 20-39%, grade 2 = 40-59%, grade 3 = 60-79% and grade 4 = 80-100%). In addition, the intensity of marker immunoreactivity was determined (grade 0 = none, grade 1 = low, grade 2 = moderate and grade 3 = strong). The product of these two grades defined the final IRS.

### Statistical analysis

Statistical data analysis was performed using PRISM 5 (GraphPad Software Inc.) on the raw data. If not otherwise specified in the figure legends, comparison of two groups in functional *in vitro* experiments was carried out using a two-sided Mann-Whitney test. If not otherwise specified in the figure legends, data are presented as dot plots with horizontal bars representing means and whiskers representing the standard error of the mean (SEM). Sample size for all *in vitro* experiments were chosen empirically. For *in vivo* experiments, sample size was predetermined using power calculations with *β*=0.8 and *α*<0.05 based on preliminary data and in compliance with the 3R system (replacement, reduction, refinement).

### Code availability

Custom code is available from the corresponding author upon reasonable request.

### Data availability

The authors declare that all data supporting the findings of this study are available within the article, its extended data files, source data or from the corresponding author upon reasonable request. Original sequencing data that support the findings of this study were deposited at the National Center for Biotechnology Information (NCBI) GEO and are accessible through the series accession number GSE120576.

## ACKNOWLEDGEMENTS

We thank Dr. E. Tomazou and Dr. N. Sheffield for help in processing of the methylation data. We thank Dr. G. Leprivier and Dr. S. von Karstedt for critical discussion, M. Melz for help in constructing TMAs, and A. Heier and A. Sendelhofert for expert technical assistance in immunohistological staining procedures. We thank Dr. V. R. Buchholz for support in cell cycle analysis. We thank Dr. D. Surdez for helpful discussion of the data. This work was mainly supported by a grant from the German Cancer Aid (DKH-111886). In addition, the laboratory of T.G.P.G. is supported by the ‘Verein zur Förderung von Wissenschaft und Forschung an der Medizinischen Fakultät der LMU München’ (WiFoMed), by LMU Munich’s Institutional Strategy LMUexcellent within the framework of the German Excellence Initiative, the ‘Mehr LEBEN für krebskranke Kinder – Bettina-Bräu-Stiftung’ (to T.G.P.G.), the Fritz-Thyssen Foundation (FTF-40.15.0.030MN), the Kind-Philipp-Foundation, the Matthias-Lackas Foundation, the Dr. Leopold and Carmen Ellinger Foundation, the Wilhelm-Sander-Foundation (2016.167.1), the German Cancer Aid (DKH-70112257), the Gert und Susanna Mayer Foundation, and the Deutsche Forschungsgemeinschaft (DFG-391665916). M.D. was supported by a scholarship of the ‘Deutsche Stiftung für junge Erwachsene mit Krebs‘, J.M. by a scholarship of the Kind-Philipp-Foundation, and T.L.B.H. by a scholarship from the German Cancer Aid. T.G.P.G., M.M. and C.A. were supported by the Swiss National Science Foundation (SNF 310030_179167). J.F.A. was supported by grants from the Cancer Prevention and Research Institute of Texas (RP120685) and the 1 Million 4 Anna Foundation.

## AUTHOR CONTRIBUTIONS

A.M. and T.G.P.G. conceived the study, wrote the paper, and drafted the figures and tables. J.S.G., M.F.O., and T.G.P.G. performed bioinformatic and statistical analyses. M.C.B. carried out gene expression analyses. A.M., M.M.L.K., J.L., and F.W. scored tissue-microarrays. S.O. and M.F.O. performed *in vivo* experiments. F.C.A., D.S., J.F.A., C.A., M.M., J.L., M.F.O., L.R.P., T.L.B.H., M.D., G.S., J.M. and S.S. contributed to experimental procedures. T.K. provided laboratory infra-structure and histological guidance. T.G.P.G. supervised the study and data analysis. All authors read and approved the final manuscript.

## COMPETING INTERESTS

The authors declare no conflict of interest.

## MATERIALS & CORRESPONDENCE

Material requests and correspondence should be addressed to Thomas G. P. Grünewald.

