## Supplementary Figures 1-6 for "Oncogenic hijacking of a developmental transcription factor evokes therapeutic vulnerability for ROS-induction in Ewing sarcoma"

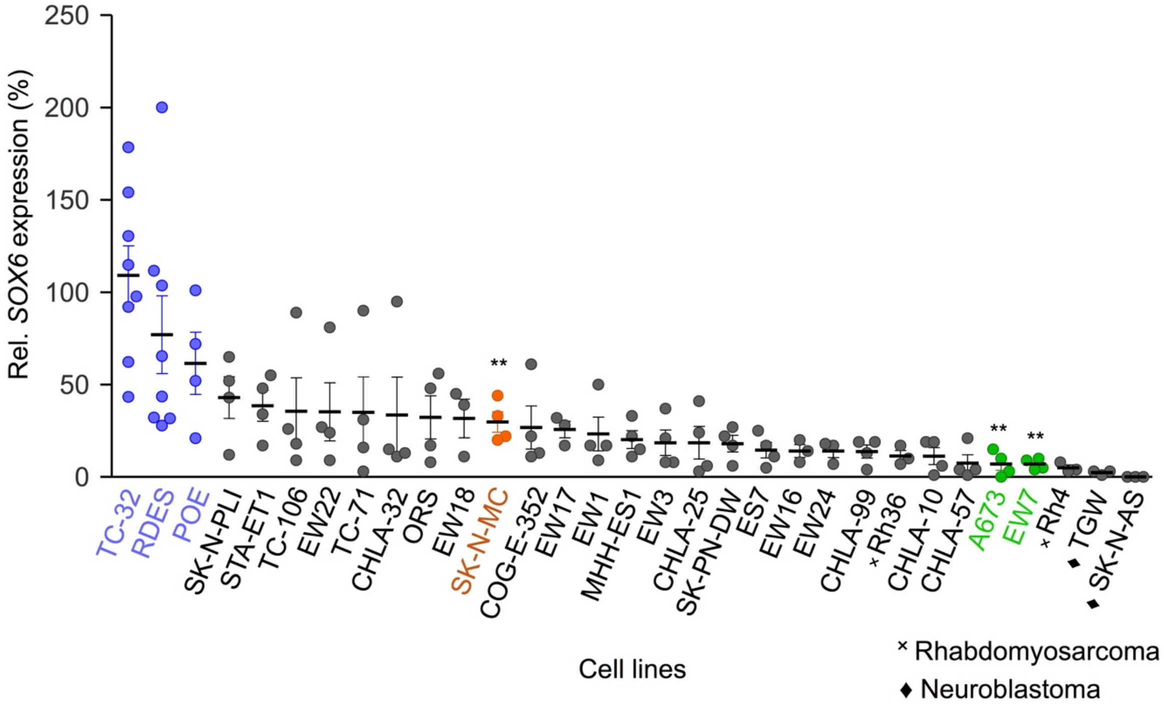


**Supplementary Fig. 1**
Analysis of *SOX6* mRNA levels by qRT-PCR of different EwS cell lines (TC-32, RDES, POE, SK-N-PLI, STA-ET1, TC-106, EW22, TC-71, CHLA-32, ORS, EW18, SK-N-MC, COG-E-352, EW17, EW1, MHH-ES1, EW3, CHLA-25, SK-PN-DW, ES7, EW16, EW24, CHLA-99, CHLA-10, CHLA-57, A673, EW7), rhabdomyosarcoma cell lines (ˣRh4, Rh36) and neuroblastoma cell lines (♦TGW, SK-N-AS). Expression levels were normalized to that of TC-32. Cell lines highlighted in blue color indicate the three EwS cell lines with highest *SOX6* expression that were used for the majority of experiments. Cell lines indicated in orange and green color represent EwS cell lines with intermediate or low *SOX6* expression, respectively. Dots represented means and whiskers SEM, $n \geq 3$.

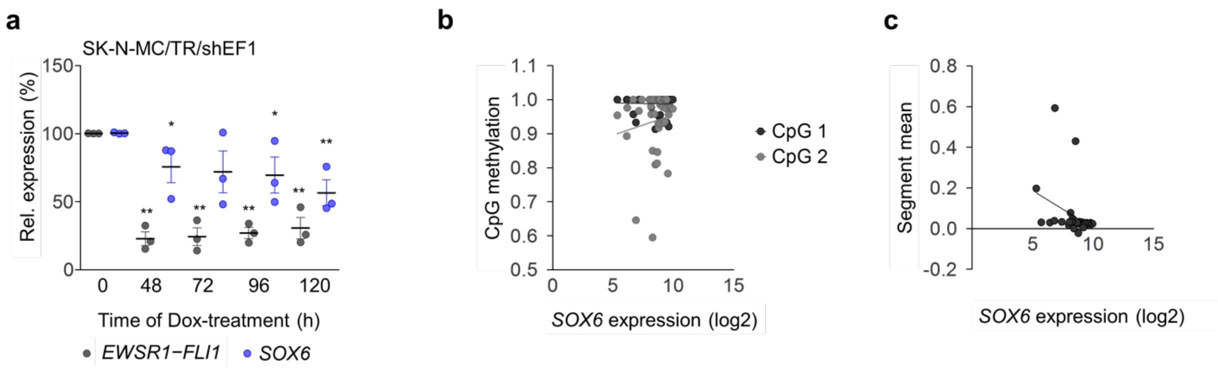


**Supplementary Fig. 2**

**a)** Analysis of *EWSR1-FLI1* and *SOX6* expression by qRT-PCR in SK-N-MC/TR/shEF1 cells at indicated time points after addition of Dox. Horizontal bars represent means and whiskers SEM, *n*=3**.** *P* values determined via two-sided Mann-Whitney test. **b)** Correlation analysis of methylation level of two CpG-methylation sites (CpG1 and CpG2) within the *SOX6* promoter and SOX6 expression levels (log2) in primary EwS tumors, *n*=40. Lines indicate linear regressions of the data. c**)** Correlation analysis of copy number variation (segment mean) at the SOX6 locus with *SOX6* expression levels (log2) in primary EwS, *n*=32. The line indicates the linear regressions of the data.

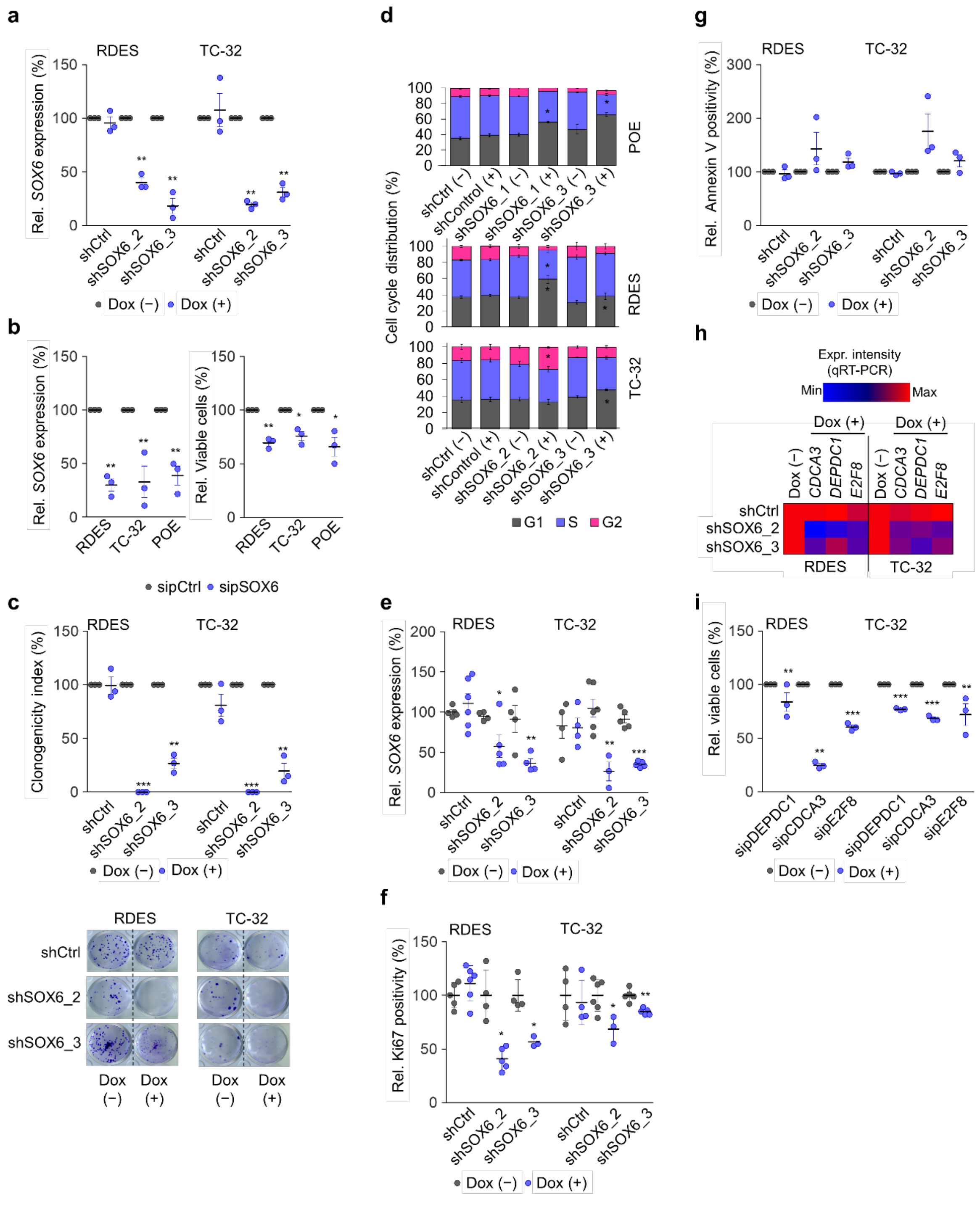


**Supplementary Fig. 3**
**a)** Analysis of relative *SOX6* expression by qRT-PCR in RDES and TC-32 EwS cell lines harboring a Dox-inducible shRNA against *SOX6* or a non-targeting control shRNA (shCtrl) 96h after addition of Dox. Horizontal bars represent means and whiskers SEM, $n$=3. $P$ values determined via two-sided Mann-Whitney t-test. **b)** Left: Analysis of relative SOX6 expression in POE, RDES and TC-32 cells transfected with a specific sipool against SOX6 or a control sipool 96h after transfection. Horizontal bars represent means and whiskers SEM, $n$=3. $P$ values determined via two-sided Mann-Whitney t-test. Right: Analysis of viable cell count of the same cells by standardized hemocytometry and Trypan blue exclusion method. Horizontal bars represent means and whiskers SEM, $n\geq3$. $P$ values determined via independent one-sample t-test. **c)** Top: Analysis of the clonogenicity index after 12 days of shRNA-mediated *SOX6*

knockdown in RDES and TC-32 cells. Horizontal bars represent means and whiskers SEM, *n*=3. *P* values determined via two-sided Mann-Whitney test. Bottom: Representative colony forming assays of the same cells. **d)** Quantification of results from flow cytometric cell cycle analysis using PI stain of POE, RDES and TC-32 EwS cells with/without shRNA-mediated *SOX6* knockdown. Cells were serum-starved for 56h and subsequently stimulated for 20h with 10% FCS. Horizontal bars represent means and whiskers SEM, *n*≥3. *P* values determined via two-sided Mann-Whitney t-test. **e)** *Ex vivo* analysis of *SOX6* expression by qRT-PCR in xenografts from RDES and TC-32 cells. Horizontal bars represent means and whiskers SEM, n≥3. **f)** *Ex vivo* analysis of Ki67 positivity of xenografted RDES and TC-32 cell lines. Horizontal bars represent means and whiskers SEM, *n*≥3. *P* values determined via two-sided Mann-Whitney t-test. **g)** Quantification of relative Annexin V positivity of RDES and TC-32 cells 96h after Dox-induced *SOX6* silencing. Horizontal bars represent means and whiskers SEM, *n*=3. **h)** Heat-map showing relative expression levels of *CDCA3*, *DEPDC1* and *E2F8* in RDES and TC-32 cells 96h after Dox-induced silencing of *SOX6*. Averaged data of *n*≥3 experiments is shown. **i)** Analysis of proliferation of RDES and TC-32 cells after sipool-mediated knockdown of either *CDCA3*, *DEPDC1* or *E2F8* for 96h. Horizontal bars represent means and whiskers SEM, *n*≥3. *P* values determined via independent one-sample t-test. ***$P$<0.001, **$P$<0.01, *$P$<0.05.

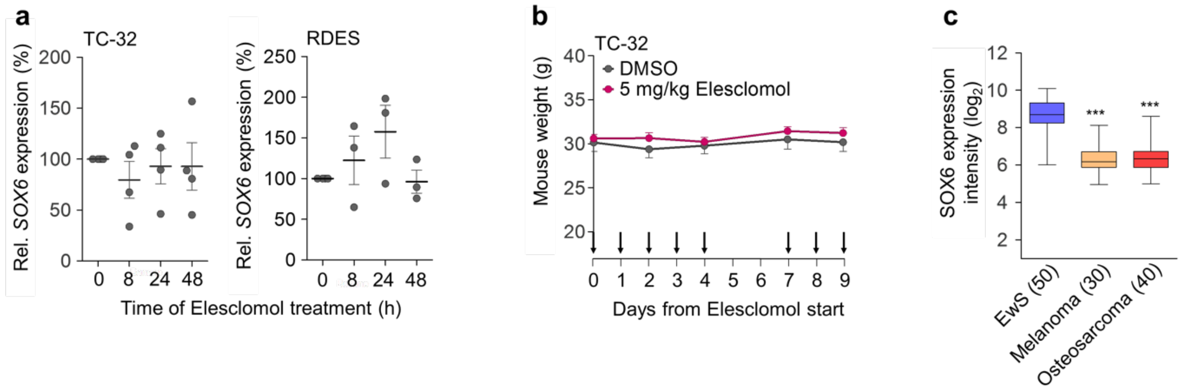


**Supplementary Fig. 4**

**a)** Quantification of relative *SOX6* expression levels in RDES and TC-32 cells at indicated time points after start of Elesclomol treatment (10 nM). Horizontal bars represent means and whiskers SEM, $n \geq 3$. *P* values determined via two-sided Mann-Whitney test. **b)** Analysis of body weight of mice over time during intravenous Elesclomol treatment (5 mg/kg). Dots represent means and whiskers SEM, $n=5$. *P* values determined via two-sided Mann-Whitney test. **c)** Analysis of *SOX6* expression intensities (log2) of primary EwS tumors, melanomas and osteosarcomas as determined by Affymetrix HG-U133Plus2.0 arrays[24]. The number of analyzed samples is given in parentheses. Horizontal bars indicate median values and boxes the interquartile range. Whiskers indicate the $10^{th}$ and $90^{th}$ percentile. *P* values determined via two-sided Mann-Whitney. ****P*<0.001.

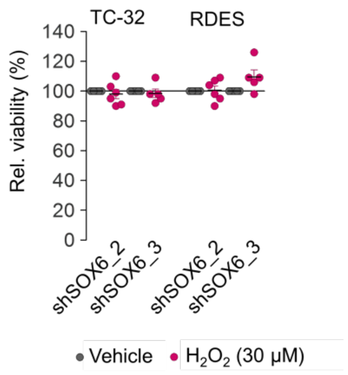


**Supplementary Fig. 5**

Analysis of cell viability of indicated cell lines treated with either vehicle ($H_2O$) or $H_2O_2$ (30 µM) by a Resazurin assay. Horizontal bars represent means and whiskers SEM, $n \geq 5$.

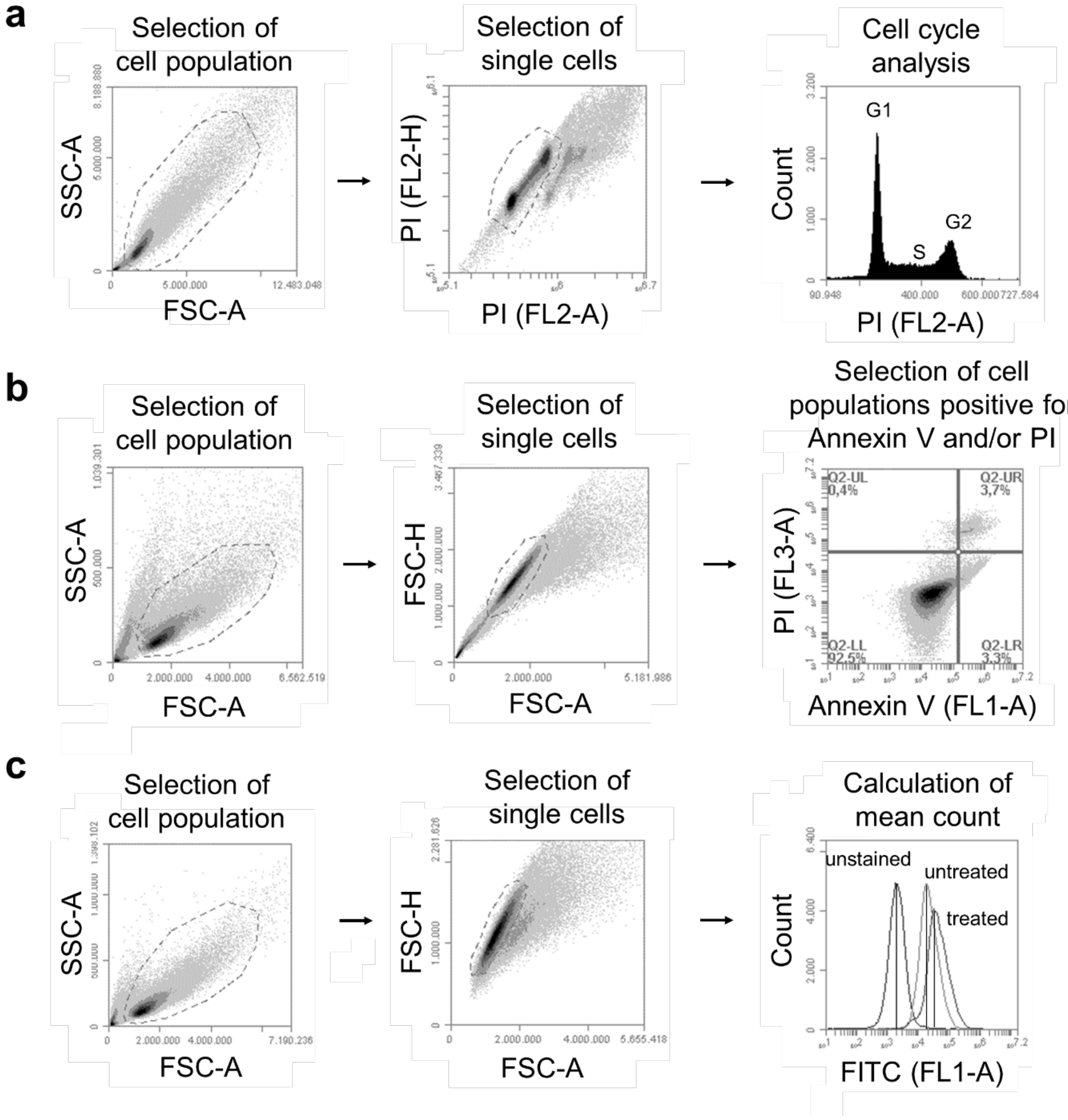


**Supplementary Fig. 6**

**a**) Gating strategy for cell-cycle analysis of EwS cells using PI. **b**) Gating strategy for Annexin V/PI staining of EwS cells. **c**) Gating strategy for ROS measurement via DCF-DA fluorescence in EwS cells treated with Elesclomol.
